# A hydrogen dependent geochemical analogue of primordial carbon and energy metabolism

**DOI:** 10.1101/682955

**Authors:** Martina Preiner, Kensuke Igarashi, Kamila B. Muchowska, Mingquan Yu, Sreejith J. Varma, Karl Kleinermanns, Masaru K. Nobu, Yoichi Kamagata, Harun Tüysüz, Joseph Moran, William F. Martin

## Abstract

Hydrogen gas, H_2_, is generated in alkaline hydrothermal vents from reactions of iron containing minerals with water during a geological process called serpentinization. It has been a source of electrons and energy since there was liquid water on the early Earth, and it fuelled early anaerobic ecosystems in the Earth’s crust^1–3^. H_2_ is the electron donor for the most ancient route of biological CO_2_ fixation, the acetyl-CoA (or Wood-Ljungdahl) pathway, which unlike any other autotrophic pathway simultaneously supplies three key requirements for life: reduced carbon in the form of acetyl groups, electrons in the form of reduced ferredoxin, and ion gradients for energy conservation in the form of ATP^4,5^. The pathway is linear, not cyclic, it releases energy rather than requiring energy input, its enzymes are replete with primordial metal cofactors^6,7^, it traces to the last universal common ancestor^8^ and abiotic, geochemical organic syntheses resembling segments of the pathway occur in hydrothermal vents today^9,10^. Laboratory simulations of the acetyl-CoA pathway’s reactions include the nonenzymatic synthesis of thioesters from CO and methylsulfide^11^, the synthesis of acetate^12^ and pyruvate^13^ from CO_2_ using native iron or external electrochemical potentials^14^ as the electron source. However, a full abiotic analogue of the acetyl-CoA pathway from H_2_ and CO_2_ as it occurs in life has not been reported to date. Here we show that three hydrothermal minerals — awaruite (Ni_3_Fe), magnetite (Fe_3_O_4_) and greigite (Fe_3_S_4_) — catalyse the fixation of CO_2_ with H_2_ at 100 °C under alkaline aqueous conditions. The product spectrum includes formate (100 mM), acetate (100 μM), pyruvate (10 μM), methanol (100 μM), and methane. With these simple catalysts, the overall exergonic reaction of the acetyl-CoA pathway is facile, shedding light on both the geochemical origin of microbial metabolism and on the nature of abiotic formate and methane synthesis in modern hydrothermal vents.

---

Organic synthesis in hydrothermal vents is relevant to life’s origin because the reactions involve sustained energy release founded in the disequilibrium between CO_2_ and the vast amounts of molecular hydrogen, H_2_, generated in the Earth’s crust during serpentinization^1,2,9,10,15–19^. Enzymatic versions of those abiotic reactions occur in core energy metabolism in acetogens and methanogens^4–7^, ancient anaerobic autotrophs that live from H_2_ and CO_2_ via the acetyl-CoA pathway and that still inhabit the crust today^7^. Though the enzymes that catalyse the microbial reactions are well investigated^4–7^, the catalysts promoting the abiotic reactions in vents today, and that might have been instrumental at life’s origin, are not fully understood^9^. To probe the mechanisms of hydrothermal metabolic reactions emulating ancient pathways, we investigated three iron minerals that naturally occur in hydrothermal vents: greigite (Fe_3_S_4_), magnetite (Fe_3_O_4_), and the nickel iron alloy awaruite (Ni_3_Fe). Although very different in structure and composition (Fig. 1), all three are geochemically synthesized in hydrothermal vents from pre-existing divalent iron and nickel minerals during serpentinization^2,16,20^. X-ray diffraction (XRD) of colloidal Fe_3_S_4_ and Ni_3_Fe nanoparticles (for details of preparation, see Methods) as well as commercial Fe_3_O_4_ reveal their characteristic pattern (Fig. 1).

**Figure 1:**
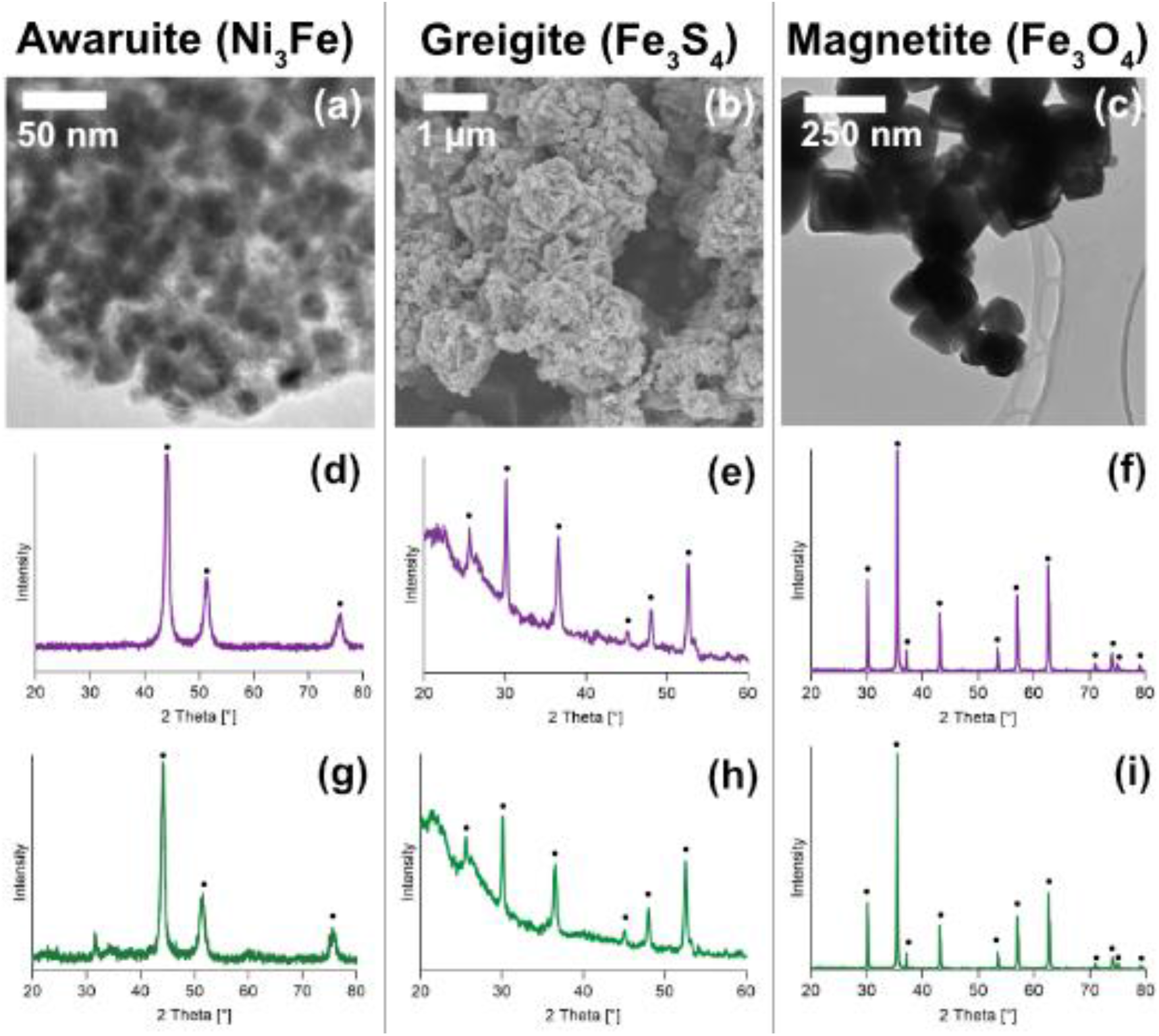
Characterization of greigite (Fe_3_S_4_), magnetite (Fe_3_O_4_) and awaruite (Ni_3_Fe) catalysts. The three powders are different in structure and composition (a, b, c), greigite and awaruite are freshly synthesized, magnetite is commercially obtained. Comparison the XRDs of the minerals before the reaction (d, e, f) and after the experiments under following conditions: Fe_3_O_4_ (h) and Ni_3_Fe (i) for 16 h under alkaline conditions (potassium hydroxide added). Fe_3_S_4_ (g), for 24 h at pH 6.5, stabilized by a phosphate buffer.

Iron sulfide minerals are formed under conditions of high H_2_S activity^16^ and have long been regarded as ancient catalysts^11,17,21^, but the key initial reaction between the inorganic and the organic world, CO_2_ fixation, has not been reported using iron sulfur catalysts under biologically relevant conditions^14^. We found that under mild hydrothermal conditions (2 bar, 100 °C) formate and acetate synthesis from H_2_ and CO_2_ occurs readily under nearly neutral and alkaline conditions in the presence of the hydrothermal mineral Fe_3_S_4_ (Fig. 2a). While only formate was detected at 20 °C, acetate also accumulates at 60 °C (Fig. 2b). At 100 bar, Fe_3_S_4_ catalyses the synthesis of formate and methane from H_2_ and CO_2_, but not from CO (Supplementary Fig. S8). Formate and methane are the main products of organic synthesis observed in the effluent of modern serpentinizing hydrothermal systems^19^. Greigite is similar in structure to the iron sulfur clusters of many modern enzymes^17^.

**Figure 2:**
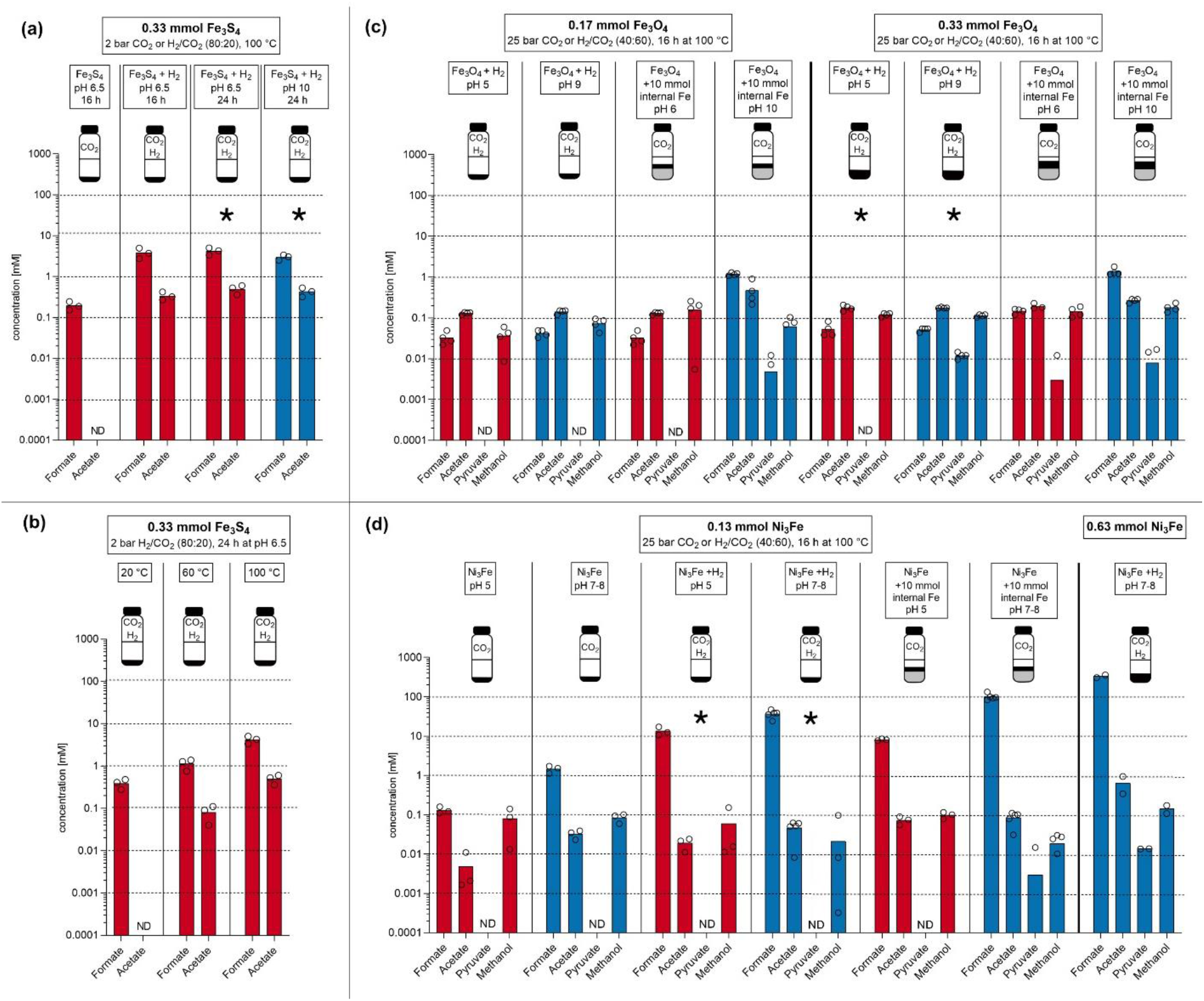
Fixation of CO_2_ with H_2_, catalysed by greigite (a,b), magnetite (c) and awaruite (d). All reactions were performed in water (1 mL for magnetite and awaruite, 3 mL for greigite). Flasks in each panel summarize the reaction parameters: hydrothermal minerals are depicted in black, additional iron powder in grey. Catalyst amounts are normalised by the number of moles of metal atoms per mole of mineral compound, 0.33 mmol of greigite (Fe_3_S_4_) or magnetite (Fe_3_O_4_), as well as 0.25 mmol of awaruite are equivalent to 1 mmol of metal atoms each (0.13 mmol awaruite are equivalent to 0.50 mmol of metal atoms). Individual experiments were performed under either CO_2_ atmosphere, H_2_/CO_2_ atmosphere, or CO_2_ atmosphere with Fe powder as electron/H_2_ source. Red bars: pH<7, Blue bars: pH>7. ND: not detected (no product was formed or product concentration was below the detection limit). Circles correspond to the values of individual experiments. Values of 0 are not shown by the logarithmic scale. Asterisks indicate experiments for which the Gibbs free energy was calculated in Tab. 1. Concentration values and standard deviations of the experiments are listed in the supplementary information (Tab.1), as are control experiments (Figs. S3-S6). The influence of pH (4–10) and reaction time (0–24 h) on the reactions catalysed by greigite is shown in supplementary material (Fig. S7).

Magnetite (Fe_3_O_4_), like H_2_, is a main end product of serpentinization, it is formed from water dependent oxidation of iron(II) silicates^22^. In chemical industry it is the catalyst of choice for diverse industrial processes including Haber-Bosch (fixation of N_2_) and Fischer-Tropsch hydrocarbon synthesis^18^. We found that magnetite, like greigite, catalyses the aqueous synthesis of formate and acetate in the range of 10 μM to 1 mM from H_2_ and CO_2_, but also methanol and pyruvate under mild (25 bar and 100 °C) hydrothermal conditions (Fig. 2c). Pyruvate is a crucial intermediate of carbon and energy metabolism in all microbes and a main product of CO_2_ fixation in autotrophs that use the acetyl-CoA pathway^4^. It accumulates at 5-10 μM in the presence of Fe_3_O_4_, but only under alkaline conditions tested, using either native Fe or H_2_ as the reductant (Fig 2c).

Awaruite (Ni_3_Fe) is an alloy of native metals and a product of serpentinization that forms at high H_2_ partial pressures and low H_2_S activities^16^. It is common in serpentinizing systems, where it is usually deposited as small grains. Its synthesis is thought to involve reduction of iron(II) and nickel(II) during phases of serpentinization in which very high H_2_ partial pressures are incurred^16^. At 100 °C, awaruite catalyses the synthesis of acetate and methanol in the 10-100 μM range at pH 5-8 whereby either the native alloy, native Fe, or H_2_ function as the reductant (Fig. 2d). At alkaline pH, with either native Fe or H_2_ as reductant, formate accumulates in the 100 mM range and pyruvate reaches 5-10 μM (Fig. 2d). In some experiments, we detected ethanol in concentrations up to over 100 μM (Fig. S10). Ni_3_Fe also catalyses CO_2_ fixation under alkaline conditions at 70 °C, and even small amounts of Ni_3_Fe are active in thermal gradients from 100 °C to 30 °C (Fig. S11), conditions similar to natural alkaline hydrothermal vents^23^. We observed trace amounts of ca. 19 ppm methane in awaruite catalysed reactions (Fig. S14 and S15), which is substantially less than an earlier report using H_2_ and CO_2_ for 1-2 weeks at 500 bar and 200-400 °C with awaruite as the catalyst^24^. Our findings show that awaruite is an effective catalyst for organic synthesis overnight from H_2_ and CO_2_ under hydrothermal conditions that, in terms of temperature and energetics, are mild enough to permit microbial growth. Of the catalysts employed, only awaruite showed minor alteration after reaction, probably due to mild oxidation (Fig. 1g-i).

Formate accumulation catalysed by awaruite reflects the near-equilibrium interconversion of H_2_-CO_2_ and formate. Under physiological conditions, the reducing power of H_2_ is insufficient to reduce CO_2_, which is why microbes studied so far that reduce CO_2_ with electrons from H_2_ employ flavin-based electron bifurcation to synthesize reduced iron sulfur clusters in ferredoxin for CO_2_ fixation^5,25^. With sufficient H_2_ and suitable inorganic catalysts, organic cofactors are not required for CO_2_ fixation to form products of the acetyl-CoA pathway (Fig. 2).

Sustained synthesis of reactive organic compounds was essential at the origin of metabolism and had to be thermodynamically favourable. Equations **1-5** show the redox reactions taking place between CO_2_ and H_2_ to form formate (**1**), methanol (**2**), acetate (**3**), pyruvate (**4**), and methane (**5**).

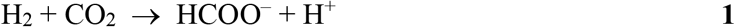

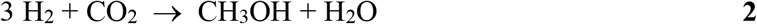

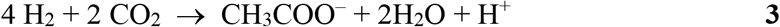

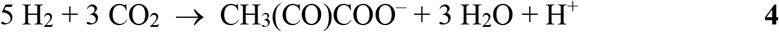

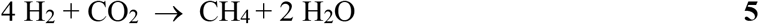

The changes in Gibbs free energy, ΔG, for six of the H_2_-dependent reactions performed here are given in Table 1 (detailed datasets are shown in supplementary Table S2). The synthesis of observed products is close to equilibrium or exergonic. For most compounds and conditions, product generation did not reach equilibrium, indicating kinetic inhibition of the reactions. Only H_2_-dependent reduction of CO_2_ to formate approached equilibrium in the presence of greigite or awaruite. Pyruvate and CH_4_ production were only detected under specific conditions despite being exergonic in nearly all treatments. For example, in treatments with H_2_ and magnetite/awaruite, pyruvate generation was only detected when the amount of mineral was increased (Fig. 2). H_2_-dependent reduction of formate to acetate (eq. 3 – eq. 1; 3H_2_ + CHOO^-^ + CO_2_ → CH_3_COO^-^ + 2H_2_O) consistently reached similar ΔG values for each mineral regardless of pH and mineral amount (roughly –70, –89, and –113 kJ mol^-1^ for greigite, magnetite, and awaruite respectively at 100 °C), suggesting the possibility of similar catalytic mechanisms. All three minerals meet kinetic inhibitions for acetate synthesis (ΔG ≪ 0) providing opportunity for energetic coupling. For those reactions in which no H_2_ was added, only native metals were available as reductant (Supplementary Figs. S5, S9a, S12, S13), likely generating intermediate H_2_ from water.

**Table 1:** Changes in Gibbs free energy ΔG for the CO_2_ fixation product formation in kJ mol^−1^.

|  | Greigite |  | Magnetite |  | Awaruite |  |
| --- | --- | --- | --- | --- | --- | --- |
| Product | 1 | 2 | 3 | 4 | 5 | 6 |
| Formate | 0.31 | -25.58 | -21.15 | -48.83 | -3.13 | -13.11 |
| Methanol | -39.57 | -39.57 | -49.44 | -47.70 | -48.75 | -53.74 |
| Acetate | -71.00 | -96.69 | -110.74 | -137.16 | -115.73 | -127.18 |
| Pyruvate | 9.07 | -15.93 | -42.90 | -71.47 | -42.90 | -57.18 |
| Methane | -152.98 | -152.98 | -187.17 | -187.17 | -187.17 | -187.17 |

**Notes:** The values of $\Delta G$ refer to the reactions as shown in equations 1-5. The conditions are those of reactions marked with asterisks in Fig. 2. Details of reaction conditions for columns 1–6: 1) 0.33 mmol Fe $_3$ S $_4$ , 100 °C, 24 h, pH 6.5, 2 bar H $_2$ /CO $_2$ (80:20); 2) 0.33 mmol Fe $_3$ S $_4$ , 100 °C, 24 h, pH 10, 2 bar H $_2$ /CO $_2$ (80:20); 3) 0.17 mmol, Fe $_3$ O $_4$ , 100 °C, 16 h, pH 6, 25 bar H $_2$ /CO $_2$ (40:60); 0.17 mmol, Fe $_3$ O $_4$ , 100 °C, 16 h, pH 9, 25 bar H $_2$ /CO $_2$ (40:60); 4) 0.17 mmol, Fe $_3$ O $_4$ , 100 °C, 16 h, pH 9, 25 bar H $_2$ /CO $_2$ (40:60); 5) 0.13 mmol, Ni $_3$ Fe, 100 °C, 16 h, pH 5, 25 bar H $_2$ /CO $_2$ (40:60); 6) 0.13 mmol, Ni $_3$ Fe, 100 °C, 16 h, pH 7-8, 25 bar H $_2$ /CO $_2$ (40:60). Columns 1, 3, 5: pH<7; Columns 2, 4, 6: pH>8.

Using greigite, magnetite or awaruite as catalysts, the synthesis of formate, acetate, methanol and pyruvate from H_2_ and CO_2_ under hydrothermal conditions is facile. The synthesis of formate and acetate is furthermore robust to the catalyst employed. The main product we observed is formate (Fig. 2), which is also the main organic product found in alkaline hydrothermal vent effluent^19^. A possible mechanism for the formation of formate is given in Figs. S18 and S19. The reaction products we observe very closely resemble those of the acetyl-CoA pathway to pyruvate^4^ (Fig. 3), which, in both the bacterial and archaeal version^4,7^, entails eleven main enzymes totalling ~15,000 amino acid residues^6^ plus six organic cofactors each with complex biosynthetic routes^7^. The bacterial and archaeal versions of the pathway involve evolutionarily unrelated enzymes but chemically similar methyl synthesis routes^4,7^. The reaction steps of the acetyl-CoA pathway employed by modern metabolism are shown in Fig. 3. The biological pathway involves the stepwise conservation of chemical energy during CO_2_ fixation as acetyl-nickel, acetyl-thioester, acetyl-phosphate, and ATP via substrate level phosphorylation (marked with an asterisk in Fig. 3).

**Figure 3:**
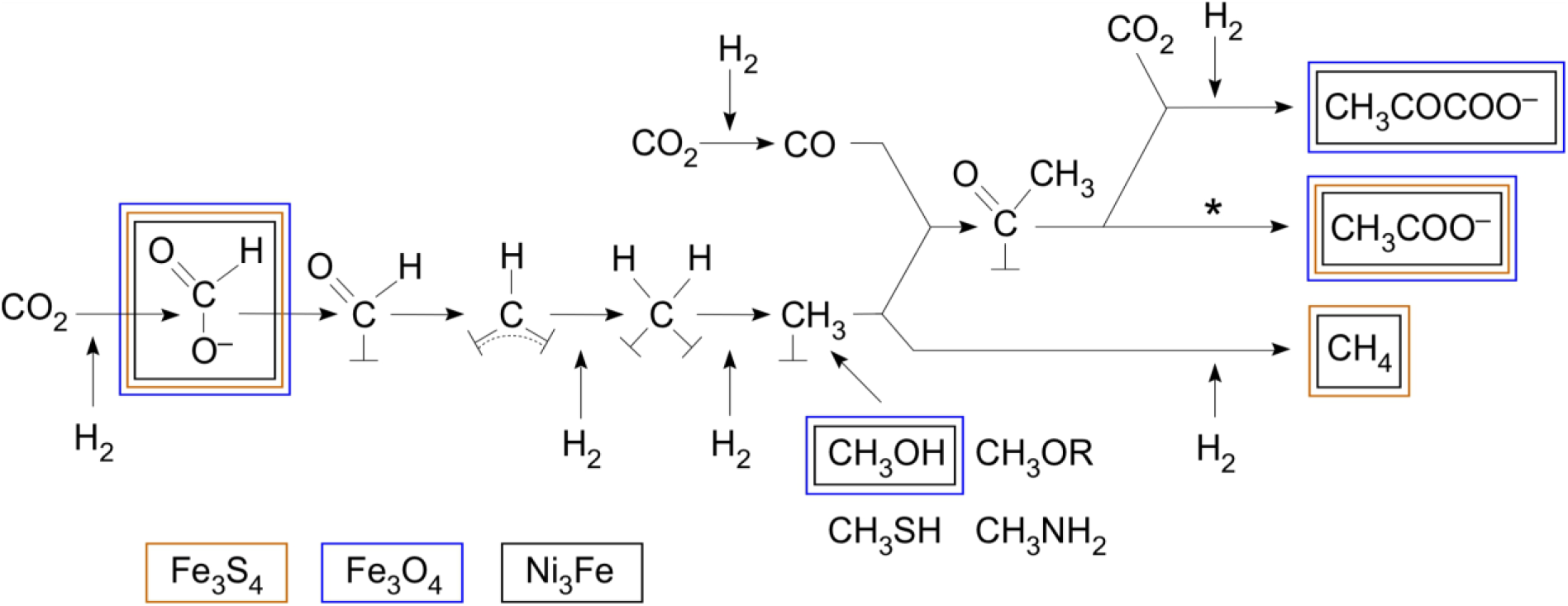
Congruence between the acetyl-CoA pathway and reactions catalysed by three iron minerals formed in hydrothermal vents. The chemical reactions summarize the acetyl-CoA pathway as it occurs in hydrogenotrophic bacteria and in archaea as depicted in ref. 4, with the exception of free formate later discovered in the archaeal pathway^26^. The methenyl (=CH–), methylene (–CH_2_–), and methyl (–CH_3_) groups of the bacterial and archaeal pathway are bound to tetrahydrofolate and tetrahydromethanopterin, respectively, generically indicated as catalysts (⊥) here. Coloured boxes indicate products observed in reactions using iron mineral catalysts. An asterisk indicates the reaction sequence in which energy is conserved as ATP via substrate level phosphorylation in the biological pathway (the acyl-nickel, thioester and acyl-phosphate intermediates that the enzymatic pathway employs for the stepwise conservation of free energy in the exergonic conversion of the nickel-bound acyl group to ATP^4^ are not shown). All products shown were observed at temperatures ≤100 °C and obtained within <24 h, except methane in the case of greigite, which was observed over the course of 24 d (Supplementary Fig. S8a). Methanol, methyl sulfide, methyl amines, and methoxy groups can serve as methyl donors for the pathway^4,27^.

Proposals for the nature of primordial CO_2_ fixation and energy conservation at biochemical origins typically posit the participation of external energy sources^28^ such as UV light^29^, heat, impact, pressure, electrical currents, or ion gradients^20^ to push organic synthesis forward. The reactions reported here require no additional energy source for a protometabolic acetyl-CoA pathway to unfold from H_2_ and CO_2_ other than the natural reactivity of two gasses and metal catalysts, indicating that membranes, though essential for the emergence of free-living cells^17^, were not required for primordial CO_2_ fixation along an exergonic, H_2_ dependent, non-enzymatic pathway to C3 products. The energy for the synthesis of compounds capable of phosphorylating ADP via substrate level phosphorylation^4,5,17^ — for reactions reported here, and for those of the enzymatically catalysed acetyl-CoA pathway, stems from the exergonic synthesis of biologically relevant organic compounds from H_2_ and CO_2_. Our findings suggest that abiotic, geochemical versions of the energy releasing reactions underlying the acetyl-CoA pathway very likely preceded the enzymes that catalyse it today^4,7,13,30^. The simplicity and primordial nature of these reactions furthermore suggest that metabolism could initiate by a similar route elsewhere, independent of light.

## Contributions

W.F.M., H.T., J.M. and M.P. designed the awaruite experiments, M.P. performed the awaruite experiments and assembled the results for the main text and SI material. K.B.M. designed and performed the magnetite experiments, S.J.V. performed exploratory experiments with magnetite. Design of the greigite experiments was done by K.I. and Y.K., K.I. performed the experiments. H.T. and M.Y. designed and synthesized the awaruite nanoparticles and performed XRD and TEM measurements for magnetite and awaruite experiments. M.K.N. performed and interpreted the thermodynamics calculations. K.K. and M.P. formulated the H_2_ reduction mechanism. W.F.M. wrote the initial draft of the main text and all authors edited the manuscript.

## Acknowledgements

We thank Yitao Dai for setting up gas analysis for the awaruite experiments, Alexander Bähr, Andrey do Nascima Vieira and Perlina Lim for scientific support and Joana C. Xavier for discussions. For funding, W.F.M. and H.T. thank the Deutsche Forschungsgemeinschaft (MA-1426/21-1 / TU 315/8-1), W.F.M thanks the European Research Council (ERC 666053) and the VW foundation (93 046). This work is partly supported by IMPRS-RECHARGE and MAXNET Energy consortium of the Max Planck Society. K.I. and Y.K. thank JSPS KAKENHI Grant-in-Aid for Scientific Research on Innovative Areas (K.I.: JP17H05240 / Y.K.: 26106004). K.I. is also supported by Grant-in-Aid for Young Scientists B (JP17K15255). J.M. thanks the European Research Council (ERC 639170) and ANR LabEX (ANR-10-LABX-0026 CSC).

## Methods

### General information

An overview of the performed experiments can be found in Fig. S1, and relevant controls – in Figs. S3-S6. The quantity of each transition metal reagent tested as carbon fixation catalyst was normalised to contain the same number of mmol of metal atoms across the experiments. For example, “1 mmol metal atoms” corresponds to: 0.33 mmol greigite Fe_3_S_4_ (99 mg), 0.33 mmol magnetite Fe_3_O_4_ (77 mg), and 0.25 mmol awaruite Ni_3_Fe (58 mg). Each reaction was performed in at least triplicate. Information on suppliers, grade and purity of all used reagents are listed in the Supplementary Information.

### Synthesis of greigite (Fe_3_S_4_)

Every piece of apparatus used in greigite synthesis was stored in an anaerobic chamber (Coy Laboratory Products) under a gas mixture of N_2_/H_2_/CO_2_ (80:5:15) for at least 48 h before use, to remove the residual oxygen. Reagents for greigite synthesis were purged with N_2_ before use unless otherwise stated. Amorphous FeO(OH) was synthesized as reported previously^31^ and suspended in Milli-Q water (0.30 mol/L) under air atmosphere. After purging with N_2_, this suspension was stored in a glass bottle under N_2_/H_2_/CO_2_ (80:5:15). The solutions of Na_2_S (1.0 M) and H_2_SO_4_ (2.0 M) were prepared as reported previously^32^ and stored in a glass bottle under N_2_. Greigite was synthesised in a solid-gas reaction system as reported previously^32^ with slight modifications. In brief, amorphous FeO(OH) (0.66 mmol, 2.2 mL of water suspension) was aliquoted to a glass reaction vessel, and a test tube containing 1.0 mL of the Na2S solution was placed in the vessel inside the anaerobic chamber. The vessel was sealed with an ETFE-coated butyl rubber stopper and an aluminium seal. Then, the vessel was removed from the anaerobic chamber and the headspace gas was replaced with Ar. After returning the vessel into the anaerobic chamber, H_2_S gas was generated inside the vessel by injecting 0.5 mL of the H_2_SO_4_ solution to the Na2S solution in the test tube using a disposable Myjector syringe (Terumo). The vessel was incubated at 80 °C for 3 hours. The resulting greigite suspension was collected by pipetting from several reaction vessels, washed with 0.5 M HCl and then rinsed with N_2_-purged Milli-Q water in the anaerobic chamber as described previously^32^.

### CO_2_ fixation catalysed by greigite

Synthesised greigite (0.33 mmol) was resuspended in 3 mL of potassium phosphate buffer (20 mM) of a designated pH. The greigite suspension was placed in a fresh glass reaction vessel, which was then sealed with an ETFE-coated butyl rubber stopper and an aluminium seal. The vessel was then removed from the chamber, and the headspace gas was replaced with H_2_/CO_2_ (80:20) or CO_2_ outside the chamber. The vessels were incubated at 100 °C over 4 to 24 h.

### HPLC analysis (greigite experiments)

Liquid phase components were analysed on a D-2000 LaChrom Elite HPLC system (Hitachi), equipped with Aminex^®^ HPX-87H column (300 mm, 7.8 mm I.D.; Bio-Rad Laboratories) and an L-2400 UV detector at 240 nm and L-2490 RI detector as described previously^33^. Supernatants obtained in the CO_2_ reduction experiments were collected after centrifugation inside the anaerobic chamber. 10 μL of the obtained supernatants were directly injected into the HPLC circuit and chromatographed under an isocratic flow of 0.7 mL/min (Eluting buffer: 10 mM H_2_SO_4_ in H_2_O). The column temperature was maintained at 50 °C. Identities of the detected analytes were determined by the LC-MS system: Agilent 1200 HPLC (Agilent Technologies) coupled to an HCT Ultra mass spectrometer (Bruker Daltonics), using a Shodex^®^ HILICpak VG-50 2D column (150 mm, 2 mm I.D.; Showa denko). The supernatant prepared as above was mixed with an equal amount of acetonitrile. Then, 5 μL of the mixture was injected into the HPLC circuit and chromatographed under an isocratic flow of 0.1 mL/min (Eluting solution: a mixture of acetonitrile and 0.25% ammonia water with 80:20 ratio). Column temperature was maintained at 30 °C.

### Synthesis of awaruite (Ni_3_Fe) nanoparticles

As previously reported^34^, spent tea leaves can be used as sustainable hard template to synthesise native metal nanoparticles in the desired composition. For the synthesis of nanoparticular Ni_3_Fe, washed and dried tea leaves were added into an aqueous solution of Ni(NO_3_)_2_·6H_2_O and Fe(NO_3_)_2_·9H_2_O (molar ratio of 3:1) and stirred at room temperature for 2 h. The mass ratio of tea leaves and metal precursors was set at 2:1. Due to the low decomposition temperature of the metal nitrate salt (below 200°C), metal oxide nanoparticles can be formed in the pore confinement of the template before its structural damage/decomposition. The carbon-based tea leaf template was burned out in air atmosphere (at 550 °C for 4 h) and the resulting Ni_3_Fe oxide was washed with 0.1 M HCl solution for 2 h and cleaned with deionized water. Finally, the product was treated in a reductive 10% H_2_/argon flow (100 mL/min) at 500 °C for 2 h to generate the intermetallic Ni_3_Fe compound^34,35^.

### CO_2_ fixation catalysed by magnetite (Fe_3_O_4_) and awaruite (Ni_3_Fe)

Awaruite and magnetite powder (commercial) were placed in a 1.5 mL glass vial. In the case of magnetite experiments, a clean PTFE-coated stir bar was added to the vial. Then, the reaction vials were filled with 1.0 mL of Milli-Q water. Whenever the effect of an increased pH of the reaction mixtures was tested, solid KOH was added into the Milli-Q water before the reaction (45 mg/mL). KOH had been tested for contaminants via the ^1^H-NMR analysis (Supplementary Fig. S16) To prevent cross-contamination while allowing for the gas to easily reach the reaction mixture, the vials were closed with caps with punctured PTFE septa. The reaction vials (3-12) were placed in a stainless-steel pressure reactor (Berghof or Parr) which was then sealed, flushed three times with ca. 5 bar CO_2_, pressurized to a final value of 25 bar CO_2_ (unless noted otherwise), and heated at the desired temperature (an external heating mantle was used) for 16 h. At a reaction temperature of 100 °C, a maximum pressure of ca. 30 bar was reached. After the reaction, the reactor was allowed to cool down to room temperature (3–4 h from 100 °C, 2–3 hfrom 70 °C) before sample analysis^13,30^.

### Experiments with iron powder or hydrogen gas

The experiments were performed according to the general procedure described above, except that 10 mmol (560 mg) Fe^0^ powder was first placed in the reaction vials, followed by the mineral tested. Further experiments exploring the impact of the amount of Fe^0^ powder are displayed in the supplementary material (Fig. S12).

Whenever H_2_ was used in the experiments, the pressure reactor was first flushed with CO_2_, then pressurized with 10 bar of H_2_ and then brought to 25 bar by adding CO_2_ again (H_2_/CO_2_ approximately 40:60).

### Work-up procedure for reaction mixtures (Ni_3_Fe and Fe_3_O_4_)

The pH of individual reaction mixtures was determined via TRITEST L pH 1–11 pH papers (Macherey-Nagel) directly after the reaction. The CO_2_ dissolved in the reaction mixture during the reaction decreased the reaction pH values due to the formation of carbonic acid. Reaction mixtures that did not contain KOH were either treated with ca. 45 mg solid KOH per 1 mL reaction mixture to precipitate the metal ions as hydroxides (in the case of Fe_3_O_4_), or left untreated (in the case of Ni_3_Fe). All samples were then centrifuged at 13.000 rpm for 10 min. The supernatant was then separated from the precipitate (catalyst) and stored at 4 °C overnight or longer until the NMR or HPLC analysis.

### NMR analysis (for awaruite and magnetite experiments)

Concentrations of formate, acetate, pyruvate and methanol (as methoxide) were determined by ^1^H-NMR, following the protocol established in Varma et al^13^. The supernatant of the centrifuged samples was therefore mixed with sodium 3-(trimethylsilyl)-1-propanesulfonate (DSS) D_2_O-solution as the internal standard (C**H_3_** peak at 0 ppm). NMR spectra were acquired on a Bruker Avance III – 600 or a Bruker Avance 300 spectrometer at 297 K, using a ZGESGP pulse program. 32 scans were acquired for each sample, the relaxation delay was set to 40 s (600 MHz) and 87 s (300 MHz), with a spectral width of 12315 ppm (600 MHz) or 11963 ppm (300 MHz). Analysis and integration were performed using MestReNova (10.0.2) software. Shifts of the measured products are depicted in Fig. S17.

### Powder X-ray diffraction (XRD)

XRD analysis was performed for pre- and post-reaction catalysts. For greigite, XRD specimen was prepared as described previously^32^. In brief, the sample was collected by centrifugation, and the obtained pellet was directly mounted as a slurry form on a silicon holder (SanyuShoko), and then sealed by using polyimide film (Nilaco Corporation) and vacuum grease (JEOL) to avoid possible desiccation and oxidation during the analysis. The specimen was analysed using a RINT2000 X-ray diffractometer (Rigaku) at room temperature for CuKα_1, 2_ radiation scanning at a step interval of 0.02° 2*θ* and a counting time of 2 seconds with a 2*θ* range from 20° to 60°, operating at an accelerating voltage of 40 kV at 30 mA. In order to prepare specimens for magnetite and awaruite experiments, the samples were collected, washed with Milli-Q water and dried under vacuum. XRD patterns of these specimens were collected at room temperature by using a theta-theta diffractometer (Stoe) in Bragg-Brentano geometry for CuKα_1, 2_ radiation scanning at a step interval of 0.04° 2*θ* and a counting time of 6 seconds with a 2*θ* range from 20° to 60°.

### Electron microscopy

Electron microscopic observation was conducted for pre-reaction catalysts to check their morphology. For greigite, a specimen for scanning electron microscopy was prepared as described previously^32^. Briefly, in the anaerobic chamber, greigite was rinsed at least three times with N_2_-purged Milli-Q water, dried at room temperature, and then mounted on an aluminium stab using carbon tape. The specimen was taken out from the anaerobic chamber, coated with platinum/palladium alloy with an ion sputter E102 (Hitachi) and observed on a JSM-6330F (JEOL) or JSM-7800F (JEOL) field-emission scanning electron microscope (FE-SEM) at an acceleration voltage of 5 kV. For magnetite and awaruite, specimens for transmission electron microscopy were prepared. The samples were collected and embedded in Spurr resin (hard mixture). Obtained resin blocks were trimmed using an EM TRIM milling system (Leica). Thin sections were cut from the resin blocks by using a microtome with a 35° diamond knife (Reichert Ultra-Cut), dispersed in Milli-Q water and transferred from the water surface on lacey carbon film-coated Cu grids (400 mesh), and observed on an S-5500 (Hitachi) transmission electron microscope at an acceleration voltage of 30 kV. Elemental mapping was conducted on the specimens prepared as above by using energy dispersive X-ray spectrometry (EDS) with a Si (Li) detector and ultrathin polymer window, operated at an acceleration voltage of 30 kV (Fig. S2).

### Thermodynamic calculations

For Gibbs free energy yield (ΔG) calculations, published values of ΔH and ΔG values were used^36,37^. The effect of temperature on the Gibbs free energy yield was calculated using the Gibbs-Helmholz equation. Equilibrium constants at different temperatures were adjusted using the van’t Hoff equation (detailed equations in supplementary information). Corrections based on non-standard pressures were estimated using partial molar volume changes of the reactions^38^. For any organic compounds that were not detected, an aqueous concentration of 0.1 μM was assumed. For CH_4_, a partial pressure of 10^-7^ atm was assumed when not detected. In reactions containing Fe^0^ as an electron donor (Tab. S2), the H_2_ concentration was estimated by assuming H_2_-dependent CO_2_ reduction to formate reached equilibrium. Final H_2_ and CO_2_ concentrations were estimated based on the measured products (subtracting 1 mol H_2_ per mole formate detected).

